# Interactions of SARS-CoV-2 infection with chronic obesity inflammation: a complex network phenomenon

**DOI:** 10.1101/2020.06.12.148577

**Authors:** Giovani Marino Favero, Luis Paulo Gomes Mascarenhas, Meirielly Furmann, Juliana Berton, Pedro Jeferson Miranda

## Abstract

Obesity is one of the biggest public health problems in the world, and its pathophysiological characteristics include chronic inflammation with an increase in various circulating inflammatory markers, such as acute inflammatory cytokines. Complications in the respiratory tract are related to bodily problems, which lead to a restriction of lung function due to reduced volume, inducing an increase in respiratory work. SARS-CoV-2 has a high potential for contamination by respiratory secretions and, therefore, obesity is one of the main risk factors for complications due to the association established between obesity, chronic inflammation and respiratory infection. The objective was to analyze the complex relationships between obesity and COVID-19 in a meta-analysis study using complex network modeling and the theoretical knockouts technique. Here, we identify and justify through a mathematical analysis the relationships between all the immunological agents added to the proposed immunological networks, considered as a simple evident interaction, relationship, influence, response, activation, based on our quantifiers. They performed the knockouts of all 52 vertices in the COVID-19 network and obesity - regardless of the environment, which would result in nonsense - and the COVID-19 infection network without considering obesity. The stationary flow vector (flow profile), for some knockouts of immunological interest in COVID-19 infections, was chosen IFNα, IL-6, IL-10, IL-17 and TNFα. This initial study pointed out the importance of chronic inflammation in the obese individual as an important factor in potentiating the disease caused by covid-19 and, in particular, the importance on IL-17.

## 1. Introduction

Obesity is one of the biggest public health problems in the world. The World Health Organization (WHO) projected for 2025 about 2.3 billion overweight adults and more than 700 million obese – with significant growth in Western Countries –, as in the United States of America (USA), the prevalence of obesity has risen from 23% to almost 40% in recent years, and in the United Kingdom, which has increased by approximately 10% in 25 years [1,2].

One of the pathophysiological characteristics of the obese is a chronic inflammation, with an increase in several circulating inflammatory markers, such as acute inflammatory cytokines, IL-6, TNF-α, CRP (C-reactive protein), haptoglobin (BULLO’ et al, 2003), IL-1b, IL-8 and IL-10. This chronic inflammatory state acts aggravating diseases such as dyslipidemia, metabolic syndrome, diabetes, hypertension, asthma and bronchitis [3].

Complications in the respiratory tract are related to body issues, which lead to a restriction of lung function due to reduced volume, unrelated effects of lung and bronchi, inducing an increase in respiratory work, expansion of torax, obstruction to airflow and decreased peripheral oxygen saturation. In addition, we have an inflammatory response that works synergistically aggravating the asthma, bronchitis and pneumonia [4-7].

In late 2019, the Severe Acute Respiratory Syndrome Corona Virus 2 (SARS-CoV-2), caused by the COVID-19 virus, have high potential for contamination through respiratory secretions causing a pandemic [8]. The pathogen identified as a new RNA involved in the beta coronavirus, similar to SARS-CoV [9-10] has caused thousands of deaths worldwide [11]. Initially, the majority of those affected by the complications shown in relation to age, being mainly elderly due to the presence of comorbidities, which is associated with infections caused by the virus, increases the lethality, in the same way observed if the obese manifest the same or greater risk [12,13].

During the H1N1 epidemic, diabetics and obese were considered to be more prone to the risk of contamination and individuals with a BMI ≥ 40 kg/m had a higher degree of complications [14]. Thus this situation is repeated again, where obese individuals demonstrate to be part of the high-risk group of complications of COVID-19 with the need for hospitalization and intensive care, probably due to the increased risk associated with the chronic diseases that obesity leads to [15].

Thereby, the ratio of the hospitalization rate among the patients identified with COVID-19 was 4.6 per 100,000 inhabitants in the USA, in which the rates were higher (13.8) among adults aged ≥ 65 years, of whom 89.3% had one or more underlying conditions, with 48.3% being obese, in addition to conditions such as hypertension, chronic lung disease, diabetes mellitus and cardiovascular disease, which raise the levels of intensive care [16].

Thus, obesity is one of the main risk factors for coronavirus complications [17], due to the association established between obesity, chronic inflammation, and respiratory infection. Considering such health problem scenario and this strong tool to analyze complex networks, the aim was analyze the complex relations of obesity and COVID-19 in a meta-analysis study using complex network modeling and the theoretical knockouts technique [18]. Specifically we shall cover the following purposes: a) To build a network of interactions between the most immediate and relevant immunological agents that participates in the COVID-19 respiratory infection; b) To made a similar network with COVID-19 respiratory infection and the immunological factors considering the chronic inflammation state observed in obese patients; c) To calculate, using the Theoretical Knockouts Method (TKM), the relative importance of such immunological agents in such phenomena in both networks and compare the results of this method in order to quantify how obesity interferes in the COVID-19 infection in a metabolic level.

## 2. Materials and Methods

### 2.1 Data Assembly

This section indicates and justifies the relations between all immunological agents (*i. e*., cells, molecules, etc) that will be added in the immunological networks proposed above. The agents are considered as vertices and their relations as directed edges in the network (*i. e*., graph). In immunological networks there is two ways for those agents to interact: increasing or decreasing, so when the A increases the activation of agent B, then an arrow will originate from A and terminate in B. On the other hand, if A decreases the activation of B, then an arrow will originate from B and terminate in A [18].

Additionally, since matter and energy must be conserved in the resulting networks, we cannot allow any vertex to be a dead-end, that is, there is no biological agent without an exit in the system (*i. e*., network). To deal with this situation, we insert in the network an origin and a terminus. All vertices that have no arrow (edge) entering it, will be connected to the origin – from the origin to these vertices. On the other hand, all vertices that have no arrow leaving it will be connected to the terminus – from these vertices to terminus. The origin and the terminus function as a section of a metabolic or an immunological phenomenon – that is, the zone of interest for analysis and knockouts. We cannot work with all possible interactions of the human body that COVID-19 and obesity influence. This is very reasonable, since only the local and most immediate have relevant and measurable impact in the phenomenon. Then, in a practical sense, both origin and terminus are considered as an environment for the purpose of our study. The environment consists in the set of less relevant interactions that we did not include in the network.

The production of IFNα, IFNAβ and IFNγ are one of the first innate responses to any viral infection. In this case, they are secreted by cells of the nasopharynx and bronchiolar mucosa. IFNα and IFNβ are secreted primarily by the host cell infected, whereas IFNγ is initially produced by macrophages and NK cells [19,20].

NK cells are the main mediators of innate immunity and, in general, expand sufficiently to eliminate viruses in 4 to 6 days. These cells are stimulated by the three Interferons. The interaction between NK cells and infected host cell promotes increased secretion of IL-1, IL-2 and IFNγ by the host cell [21].

The viral RNA itself functions as a Pathogen-associated molecular patterns (PAMPS), stimulating, among other cells, macrophages that, in addition to phagocytic activities, will secrete a vast amount of substances, among them: IL-1, IL-18 (which will increase secretion of IFNγ), IL-6, IL-12, TNF, NO, IL-10, IL-8. The association of IL-18 with IL-12 will promote a decrease in IL-4, which will lead to less IgE and IgG1. IL-8 will stimulate the phagocytic activity of Neutrophils associated with the complement C3b, which will cause the secretion of Myeloperoxidases, Defensin, Neutrophil elastase, Bacterial / permeability-increasing protein (BPI), Cathepsin, Lactoferrin and Gelatinases [22,23].

Dendritic cells (DC) stimulated by the viral particle and interferons will activate CD4 + T cells. The interaction between DC and CD4 + T (CD40 - CD40L) will activate Tc (increased secretion of IFNγ, IL-10), B cells (secretion of TGF-β, Th17, IL-17 and IL-10) and DC licensing. Being an important step in the progression of the response to the viral agent [24,25].

Chronic inflammation in the obese is characterized by the constant circulation of TNFα, IL-1b, IL-6, IL-8, TGF-β (leptin), IL-17 and IL-18, secreted by adipose tissue [26].

All the above stated interactions can be observed in the form of a complex network in Fig. 1.

**Figure 1.**
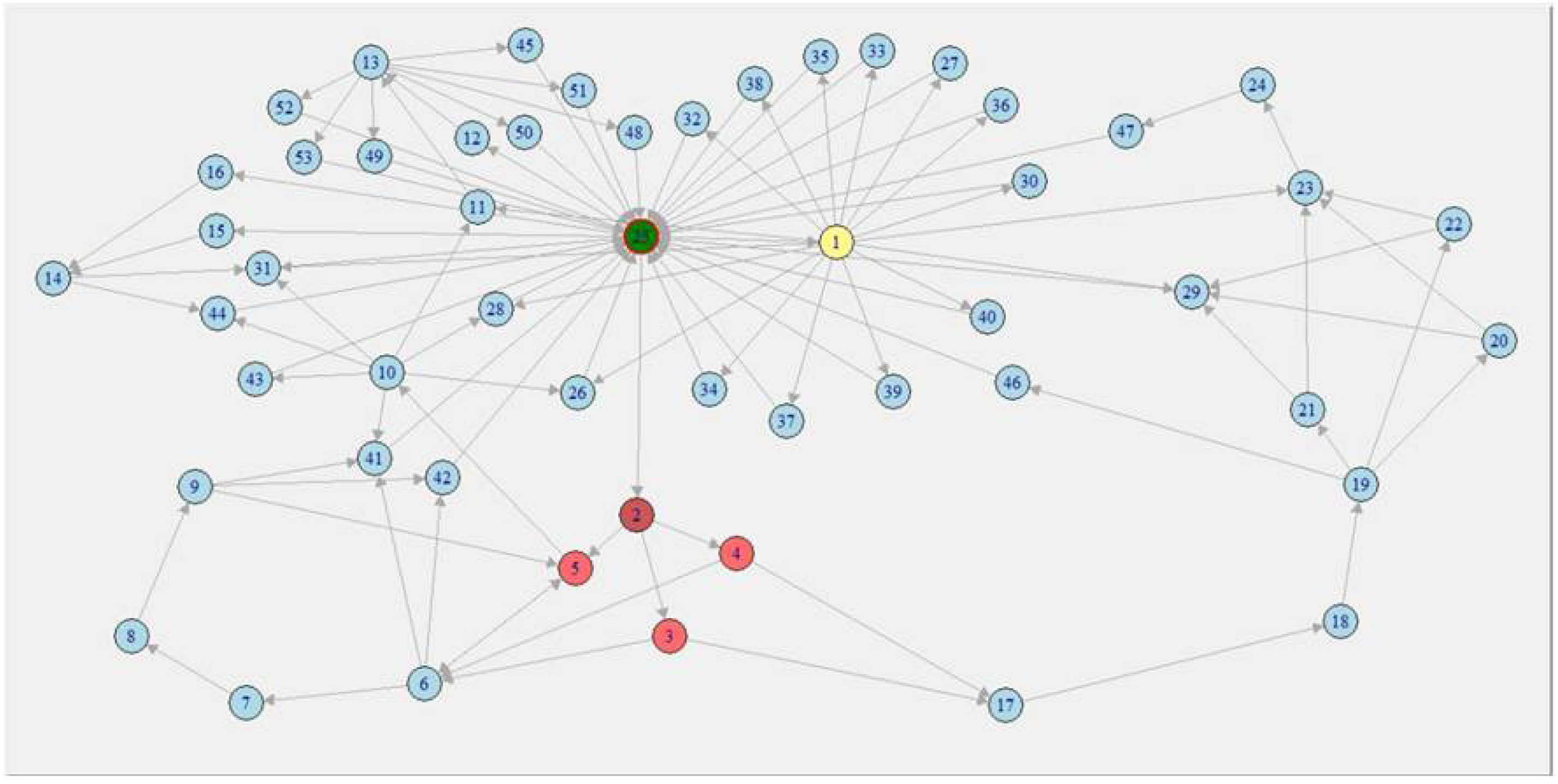
The network of immunological interactions for COVID-19 respiratory infection and chronic inflammatory state induced by obesity. Vertices labels: 1 – Adipose Tissue; 2 – COVID-19; 3 –IFNα; 4 – IFNβ; 5 – IFNγ; 6 – NK Cell; 7 – AR; 8 – AL; 9 – Infected Host Cell; 10 – Macrophage; 11 – IL-8; 12 – C3b; 13 – Neutrophil; 14 – IL-4; 15 – IgE; 16 – IgG1; 17 – DC; 18 – CD40; 19 – CD40L; 20 – DC licensing; 21 – Tc activation; 22 – B cell activation; 23 – TGF-β; 24 – Th17; 25 – Environment; 26 – TNFa; 27 – IL1b; 28 – IL-6; 29 – IL-10; 30 – IL-17D; 31 – Il-18; 32 – PAI-1; 33 – Heptoglobin; 34 – Serum Amyloid A; 35 – α1-Acid glycoprotein; 36 – 23p3; 37 – CRP; 38 – Adiponectin; 39 – NGF; 40 – MCP-1; 41 – IL1; 42 – IL-2; 43 – NO; 44 – IL-12; 45 – Myeloperoxidase; 46 – CD4*T; 47 – IL-17; 48 – Defensins; 49 – Neutrophil elastase; 50 – Bactericidal/BPI; 51 – Cathepsin; 52 – Lactoferrin; 53 – Gelatinase.

### 2.2. Summarization of the theoretical knockouts’ theory and technique

The relations (edges) must be considered as a simple evident interaction, relation, influence, response, activation.

After the building of the network – which is, for our purposes, synonym of graph. A graph is an ordered pair *G* = (*V,E*) in which *V* is a set of vertices and *E* is a subset of *V* composed by edges. The graph obtained by the above stated method is connected. This means that given any pair of vertices A and B of a graph *G*, there is always at least one directed path between them – a set of directed edges from A to B or from B to A. This condition is mandatory for the calculations presented bellow. Furthermore, such condition guarantees that all matter and energy that flows in the network is conserved, which is a very reasonable condition.

Mathematically, we do not work with the graph object as it is designed in Figure 1, but we work with another way to represent it: the adjacency matrix of a graph. This matrix is build with the following consideration: if there is a directed edge (arrow) connecting the vertices *i* and *j*, then the value of the adjacency matrix’s element *a*_*ij*_ equals 1, and otherwise equals 0. In order to calculate the distribution of probabilities, we must define the out-degree of a vertex. Let *x* be a vertex of the graph, the out-degree of *x* is the number of edges that originates in *x*. Mathematically, and using the concept of adjacency matrix, we have: the out-degree of vertex *x* is 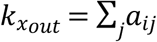.

All vertices in the network interact dynamically, that is: every vertex generates a signal of increase or decrease of biological activity. To model this dynamics, we used the random walk in the network. The time variable is added to the network, when times increases (discrete), a walker is created in the vertex environment; then if *t* is time, the total number of walkers in the network is *N*(*t*) = *t*. The walkers (*i. e*., particles, information, stimuli, activation, etc) transits in the network from vertex to vertex, one step per walker per unit of time. The amount (discrete) of walkers in vertex *i* in the time *t*isdenoted by *σ*_*i*_(*t*). Ergo, the relative number of walkers – for now on, information – in a vertex *i* is coined as local flux, defined as 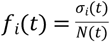. Naturally, as times evolves, the values of *f*_*i*_(*t*) for any *i*vertex in the network changes. If there are *n* vertices in the network, then there will be *n* values of *f*_*i*_ for every time *t*. Since we want to study the general state of the network, we devised a state vector coined as Flux Vector given by *F*_*V*(*G*)_(*t*) = (*f*_1_(*t*), *f*_2_(*t*), *f*_3_(*t*),…,*f*_*n* – 1_(*t*), *f*_*n*_(*t*)).

Thus, for every time *t*, there is a flux vector *F*_*V*(*G*)_(*t*). However, we are interest in generating a steady measurement, for this we want to compute the stationary state of the dynamics performed upon the network. The stationary state of the flux vector for a given network is a vector that does not change with the increase of time, denoted by *F*_*V*(*G*)_(*t*→∞) = (*f*_1_’(*t*→∞), *f*_2_’(*t*→∞), *f*_3_’(*t*→∞),…,*f*_*n* – 1_’(*t*→∞), *f*_*n*_’(*t*→∞)). Taking into account the stationary state *F*_*V*(*G*)_(*t*→∞). In dynamics, there is three possible states when times increases indefinitely: a) or it is periodic, that is: the vector *F*_*V*(*G*)_ transits within a set of states with a certain period; b) or it is chaotic, that is: the vector *F*_*V*(*G*)_ transits through a set of infinite states, never repeating; c) or it is stationary, there is only one state as times increases.

In order to test these possible options, it is needed to define the transition matrix *T* of a given graph *G*. This matrix is also known as probability matrix, because its entries are probabilities given by 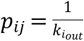. This matrix has a algebraic interest in our study, since it can be used for model time, algebraically: *T*.*F*_*V*(*G*)_(*t*) = *F*_*V*(*G*)_(*t* + 1). Every time the transition matrix operates upon *F*_*V*(*G*)_(*t*), times goes on in the dynamics of the flux in the network. Equivalently: *T*^*t*^.*F*_*V*(*G*)_(*t* = 0) = *F*_*V*(*G*)_(*t*)⇒*T*^*t*→∞^.*F*_*V*(*G*)_(*t* = 0) = *F*_*V*(*G*)_(*t*→∞). It is possible to compute numerically *T*^*t*→∞^ as a limit, but we are interested in an exact evaluation of *F*_*V*(*G*)_(*t*→∞). For this, we must consider the Perron-Frobenius features of the transition matrix *T*. These features allows one to assert that there exists an unique stationary state *F*_*V*(*G*)_(*t*→∞), and it can be computed exactly by normalizing the eigenvector associated to the major eigenvalue of *T*. It wasconsidering the set {λ_*i*_} of eigenvalues of *T*. If the matrix *T* met the following criteria, possessing the Perron-Frobenius features: a) |λ_1_| ≥ |λ_2_| ≥ |λ_3_| ≥ …|λ_*n* – 1_| ≥ |λ_*n*_| and b) |λ_1_| = 1.

In our concrete case, since the graph is connected – due to the conservation of matter and energy in the time –, it also met the Perron-Frobenius features. This means that *F*_*V*(*G*)_(*t*→∞) can be calculated exactly, with no need for numeric computation, by normalizing the eigenvector associated to the biggest eigenvalue: 1.

An initial graph*G* represents the normal functioning of a phenomenon of interest. When a knockout to a given vertex *i*from *G*, generates in this process a knocked-out graph *G’*. Considering that the transition matrix *T* of *G* met the Perron-Frobenius features, then we can compute the stationary state *F*_*V*(*G*)_(*t*→∞). Since *G’* is derived from *G*, it will be also connected, ergo possessing the Perron-Frobenius features. This means that we can also compute its stationary state *F*_*V*(*G*’)_(*t*→∞). Using both *F*_*V*(*G*’)_ and *F*_*V*(*G*)_ one can calculate the distance between such vectors, given by: *D*_*G,G*’_ = *F*_*V*(*G*)_(*t*→∞) – *F*_*V*(*G*’)_(*t*→∞) = (Δ*f*_1_,Δ*f*_2_,Δ*f*_3_,…,Δ*f*_*n* – 1_,Δ*f*_*n*_ = *f*_*n*_)., where 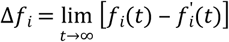 for 1 ≤ *i* ≤ *n*. Naturally, the values of Δ*f*_*i*_ can be positive or negative; if it is positive, it means that the activation of the component is locally increase by the knockout (KO); however, if it is negative, it means that the activation is locally decreased. For each Δ*f*_*i*_, we can compute its relative error, by:

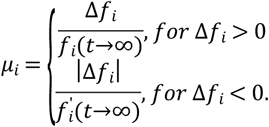

Collecting all relative error, we can average the set of all error and obtain the relative mean error, which is a global measure: 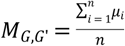. This measure is an index, that is, ranges from 0 to 1. If *M*_*G,G*’_ is close to 0, then the particular KO was not significant to the normal functioning of the graph *G*. On the other hand, if *M*_*G,G*’_ is close to 1, then the KO is very relevant to the normal functioning of the graph *G*.

It is important to note that our main measures are: the stationary state *F*_*V*(*G*’)_(*t*→∞) due to a particular KO (also known as Flux Profile): it shows how the local fluxes change with particular KO; and the relative mean error *M*_*G,G*’_, which indicates how much a local KO can impact globally. Both quantifiers are very steady and can easily be biologically interpreted in a variety of biological phenomena. Now, based on our quantifiers we performed the knockouts of all 52 vertices in the COVID-19 and obesity network – regardless of the environment, which would result in nonsense -, and the COVID-19 infection network not considering obesity.

## 3. Results

### 3.1 Relative Mean Error measurements

Since we built two networks, there are two sets of results concerning the Relative Mean Error (RME). The first network models the COVID-19 and obesity happening together. And the second one models only the effects of COVID-19, Network 1 and Network 2, respectively. Table 1 shows the values of RME and its deviations for both networks. In the sequel, we have plotted a histogram (Figure 2) of these.

**Table 1.** Summarization of relative mean errors (RME) and its deviations to Network 1 and Network 2. The values are decreasing for both cases.

| Immunological Agent<br>(Network 1) | RME<br>(Network 1) | Deviation | Immunological Agent<br>(Network 2) | RME<br>(Network 2) | Deviation |
| --- | --- | --- | --- | --- | --- |
| Infected Host Cell | 0.9010 | $\pm 0.1870$ | COVID | 1.0000 | $\pm 0.0000$ |
| COVID | 0.8645 | $\pm 0.2407$ | Infected Host Cell | 0.8289 | $\pm 0.2618$ |
| CD40L | 0.8559 | $\pm 0.2752$ | CD40L | 0.8151 | $\pm 0.3131$ |
| IL-17 | 0.8260 | $\pm 0.3265$ | IL-17 | 0.7817 | $\pm 0.3457$ |
| CD40 | 0.7929 | $\pm 0.3227$ | CD40 | 0.7807 | $\pm 0.3214$ |
| AR | 0.7857 | $\pm 0.2586$ | DC | 0.7642 | $\pm 0.3238$ |
| Adipocyte | 0.7644 | $\pm 0.2918$ | Neutrophil | 0.6932 | $\pm 0.3346$ |
| AL | 0.7634 | $\pm 0.3008$ | NK Cell | 0.6878 | $\pm 0.2731$ |
| Th17 | 0.7540 | $\pm 0.3183$ | AL | 0.6819 | $\pm 0.3053$ |
| Neutrophil | 0.7509 | $\pm 0.3309$ | IL-4 | 0.6627 | $\pm 0.3477$ |
| IL-4 | 0.7304 | $\pm 0.3565$ | AR | 0.6505 | $\pm 0.2886$ |
| Macrophage | 0.7286 | $\pm 0.2828$ | IFNα | 0.6493 | $\pm 0.2915$ |
| IFN-γ | 0.7156 | $\pm 0.3044$ | IFNβ | 0.6493 | $\pm 0.2915$ |
| NK Cell | 0.6875 | $\pm 0.3197$ | IFNγ | 0.6289 | $\pm 0.3081$ |
| DC | 0.6807 | $\pm 0.3469$ | Th17 | 0.6214 | $\pm 0.3273$ |
| IL-8 | 0.6371 | $\pm 0.2650$ | IL-8 | 0.6099 | $\pm 0.2586$ |
| TGF-β | 0.6369 | $\pm 0.3371$ | C3b | 0.5945 | $\pm 0.3215$ |
| IFNα | 0.6351 | $\pm 0.2870$ | Tc activation | 0.5796 | $\pm 0.2735$ |
| Tc activation | 0.6351 | $\pm 0.3072$ | B cell activation | 0.5796 | $\pm 0.2735$ |
| B cell activation | 0.6351 | $\pm 0.3072$ | DC licensing | 0.5736 | $\pm 0.2678$ |
| DC licensing | 0.6351 | $\pm 0.3072$ | Macrophage | 0.5494 | $\pm 0.3168$ |
| IFNβ | 0.6351 | $\pm 0.2871$ | IgE | 0.5370 | $\pm 0.3471$ |
| IL-1 | 0.6167 | $\pm 0.3786$ | IgG1 | 0.5370 | $\pm 0.3471$ |
| IgE | 0.5948 | $\pm 0.3631$ | TGF-β | 0.5094 | $\pm 0.3286$ |
| IgG1 | 0.5842 | $\pm 0.3592$ | NO | 0.5049 | $\pm 0.4036$ |
| C3b | 0.5673 | $\pm 0.3508$ | IL-2 | 0.4967 | $\pm 0.3751$ |
| IL-18 | 0.5420 | $\pm 0.3314$ | Defensins | 0.4948 | $\pm 0.4204$ |
| Heptoglobin | 0.5203 | $\pm 0.3767$ | Neutrophil elastase | 0.4919 | $\pm 0.4173$ |
| Serum amyloid A | 0.5203 | $\pm 0.3767$ | Bactericidal/BPI | 0.4918 | $\pm 0.4172$ |
| PAI-1 | 0.5163 | $\pm 0.3749$ | Cathepsin | 0.4918 | $\pm 0.4172$ |
| 23p3 | 0.5138 | $\pm 0.3753$ | Lactoferrin | 0.4918 | $\pm 0.4172$ |
| Adiponectin | 0.5130 | $\pm 0.3747$ | Gelatinases | 0.4918 | $\pm 0.4172$ |
| NGF | 0.5130 | $\pm 0.3747$ | IL-12 | 0.4848 | $\pm 0.3744$ |
| MCP-1 | 0.5130 | $\pm 0.3747$ | IL-18 | 0.4827 | $\pm 0.3514$ |
| CRP | 0.5128 | $\pm 0.3745$ | TNFα | 0.4648 | $\pm 0.4119$ |
| α1-Acid glycoprotein | 0.5128 | $\pm 0.3746$ | IL-6 | 0.4594 | $\pm 0.4141$ |
| IL-17D | 0.4291 | $\pm 0.3879$ | Myeloperoxidase | 0.4555 | $\pm 0.4165$ |
| NO | 0.4246 | $\pm 0.3564$ | IL-1 | 0.4505 | $\pm 0.3752$ |
| IL-1b | 0.4245 | $\pm 0.3895$ | IL-10 | 0.4222 | $\pm 0.3294$ |
| CD4*T | 0.4240 | $\pm 0.3667$ | CD4*T | 0.3939 | $\pm 0.3923$ |
| IL-12 | 0.4162 | $\pm 0.3039$ | Adipocyte | 0.0000 | $\pm 0.0000$ |
| Myeloperoxidase | 0.4132 | $\pm 0.3708$ | Heptoglobin | 0.0000 | $\pm 0.0000$ |
| IL-2 | 0.4052 | $\pm 0.3187$ | Serum amyloid A | 0.0000 | $\pm 0.0000$ |
| Defensins | 0.3992 | $\pm 0.3407$ | PAI-1 | 0.0000 | $\pm 0.0000$ |
| Neutrophil elastase | 0.3990 | $\pm 0.3402$ | 23p3 | 0.0000 | $\pm 0.0000$ |
| Bactericidal/BPI | 0.3990 | $\pm 0.3402$ | Adiponectin | 0.0000 | $\pm 0.0000$ |
| Cathepsin | 0.3990 | ±0.3402 | NGF | 0.0000 | ±0.0000 |
| Lactoferrin | 0.3990 | ±0.3402 | <u>MCP-1</u> | 0.0000 | ±0.0000 |
| Gelatinases | 0.3990 | ±0.3402 | CRP | 0.0000 | ±0.0000 |
| TNFα | 0.3965 | ±0.3501 | <u>α1-Acid glycoprotein</u> | 0.0000 | ±0.0000 |
| IL-6 | 0.3849 | ±0.3394 | <u>IL-17D</u> | 0.0000 | ±0.0000 |
| IL-10 | 0.2994 | ±0.3097 | <u>IL-1b</u> | 0.0000 | ±0.0000 |
| *Environment* | 0.0000 | ±0.0000 | *Environment* | 0.0000 | ±0.0000 |

**Figure 2.**
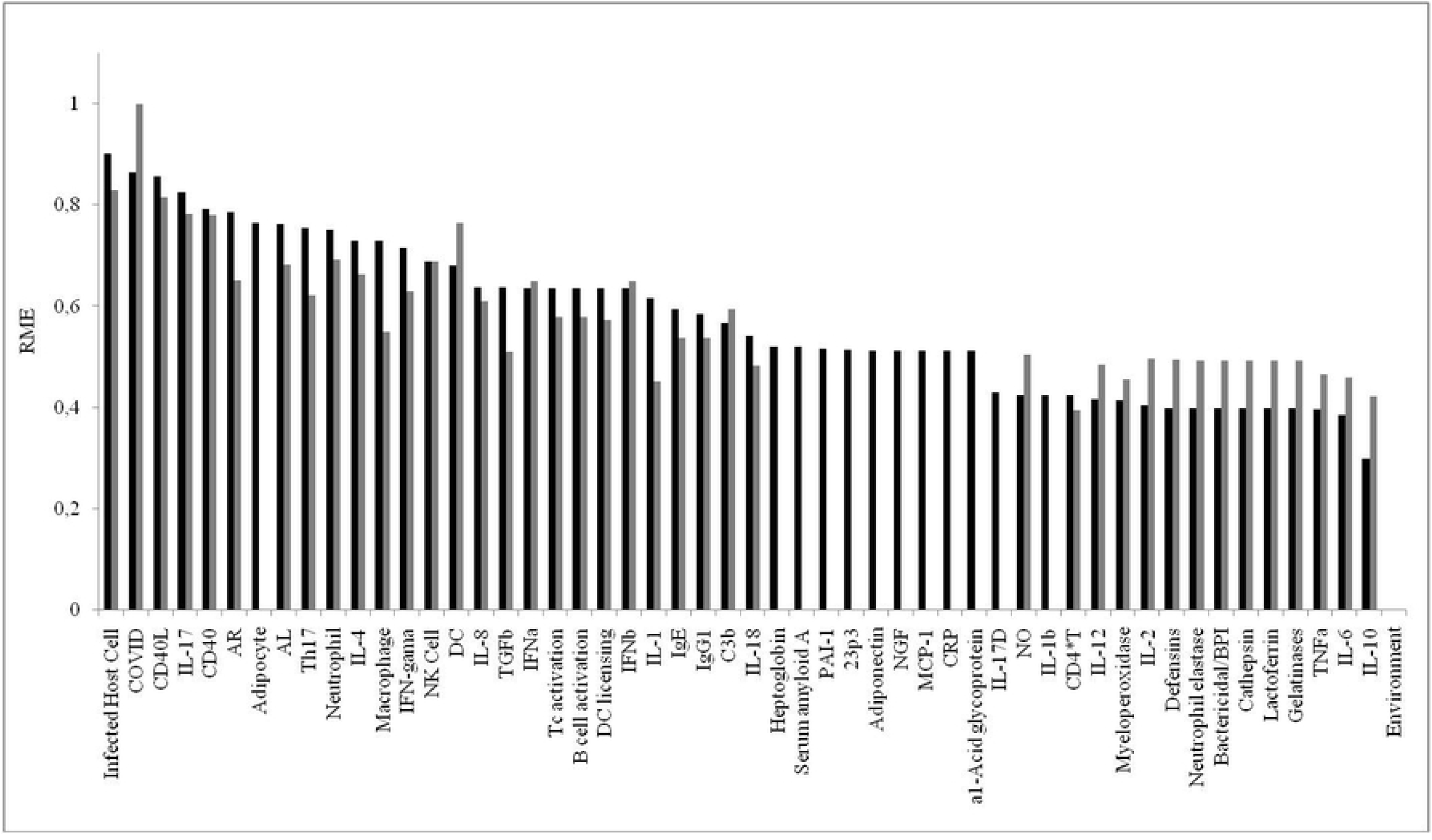
Histogram of the RME of knockouts from Network 1 (black) and Network 2 (grey).

### 3.2 Flux profiles

In this section, we show the Stationary Flux Vector (Flux Profile) for some knock-outs of immunological interest in COVID-19 infections. It was chosen IFNα, IL-6, IL-10, IL-17, and TNFα (Figures 3-6, respectively) as focus for discussion. Other knock-outs are presented as supplementary material of the present paper, which are: IFNβ,IFNγ, IL-1, IL2, IL-4, IL-8, and IL-18. It is important to state that these Flux Profiles have as base the Network 1 (COVID-19 and obesity).

**Figure 3.**
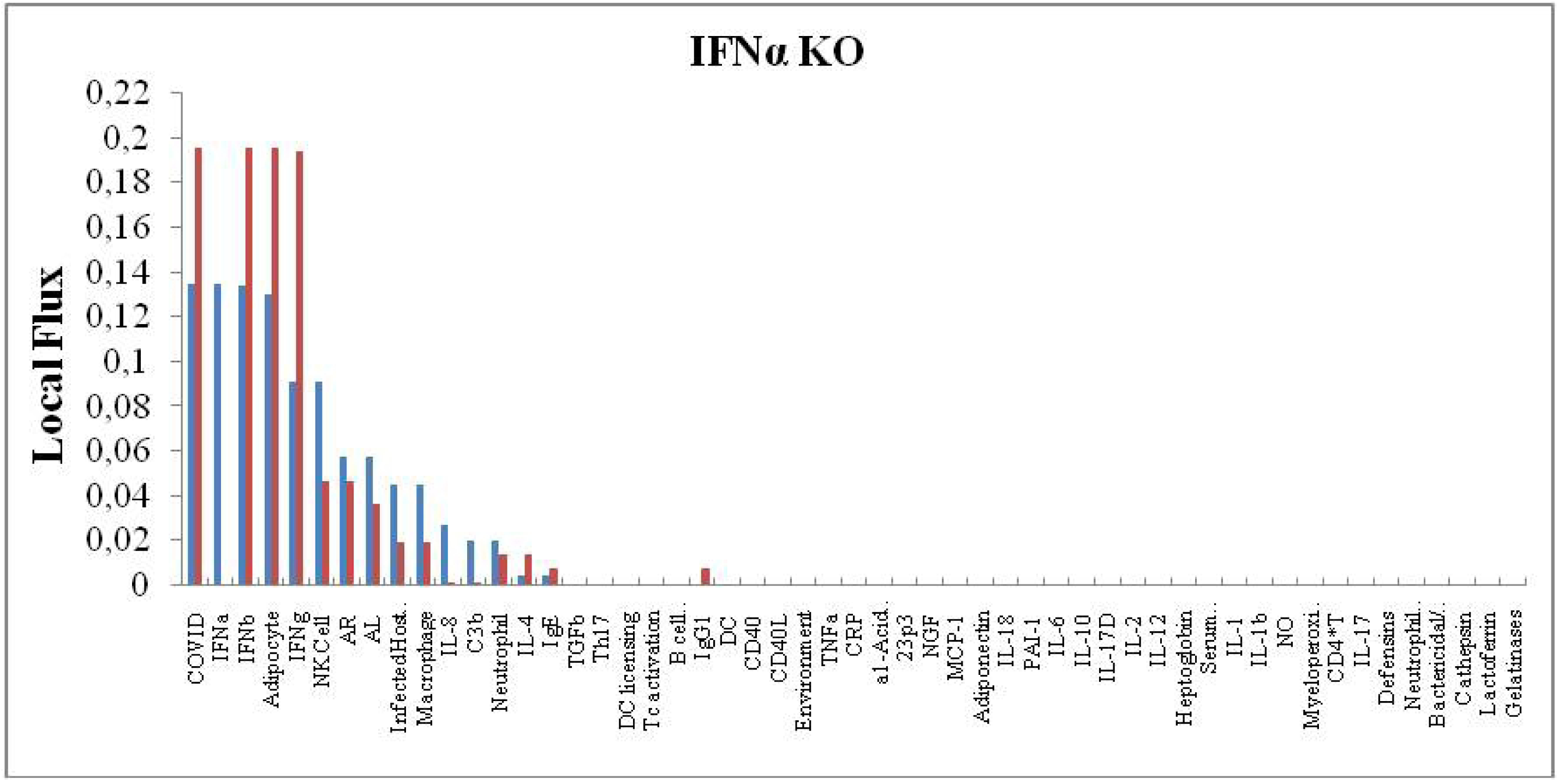
Flux Profile for IFNα knockout and the effect on local fluxes. The blue histogram stands for the Standard Flux (without KO) and the red histogram stands for the KO flux.

**Figure 4.**
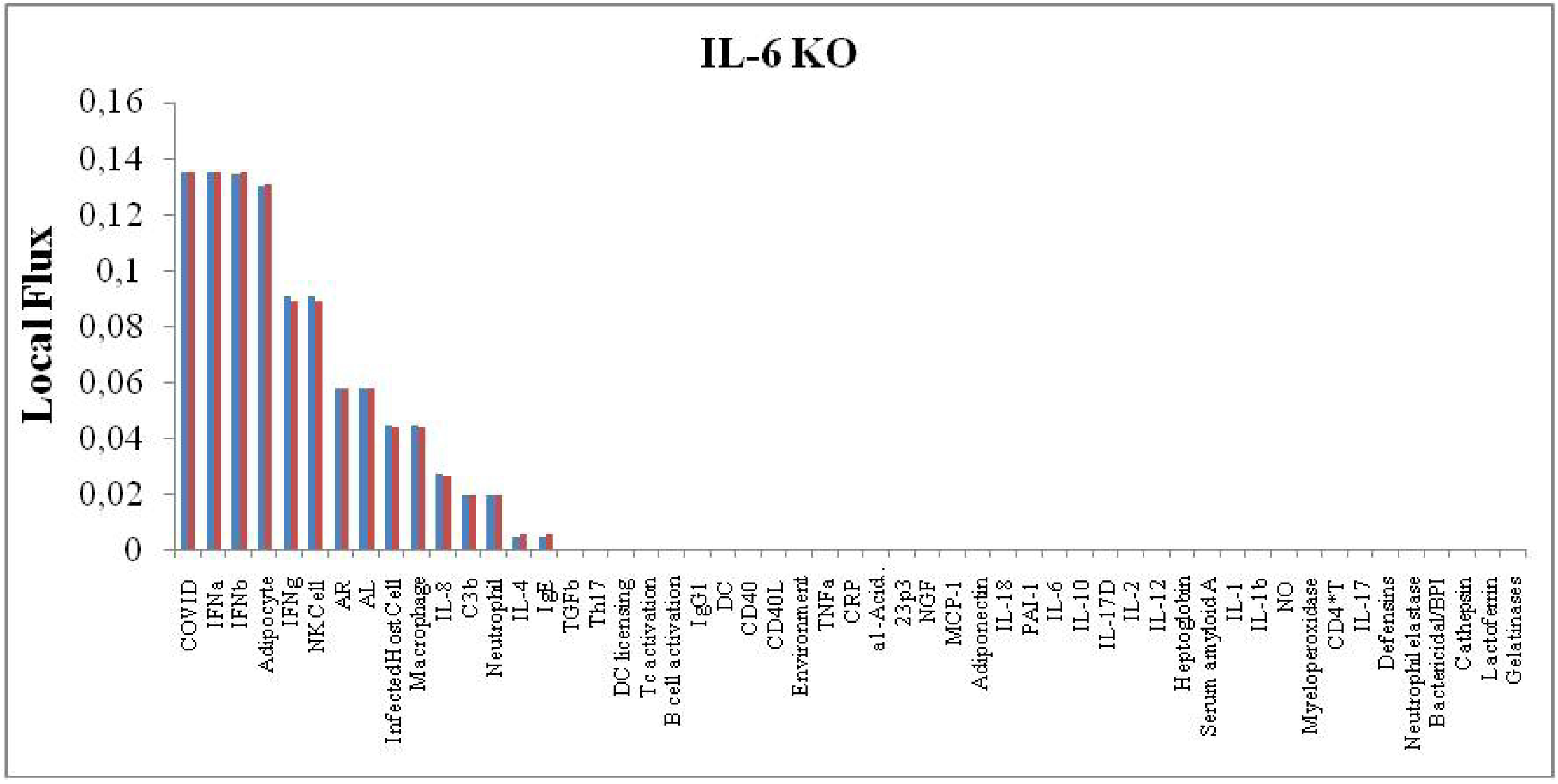
Flux Profile for IL-6 knockout and the effect on local fluxes. The blue histogram stands for the Standard Flux (without KO) and the red histogram stands for the KO flux.

**Figure 5.**
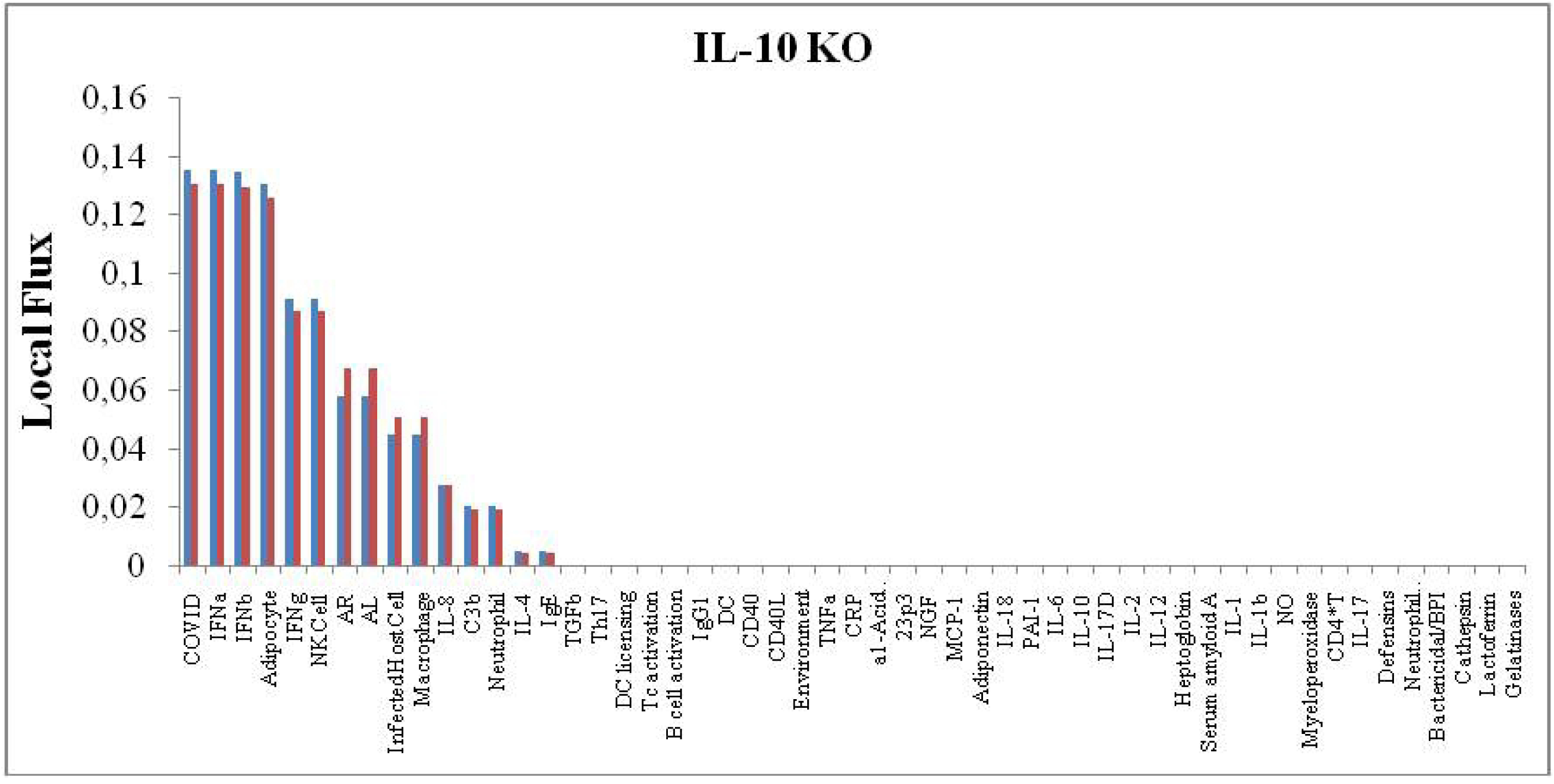
Flux Profile for IL-10 knockout and the effect on local fluxes. The blue histogram stands for the Standard Flux (without KO) and the red histogram stands for the KO flux.

**Figure 6.**
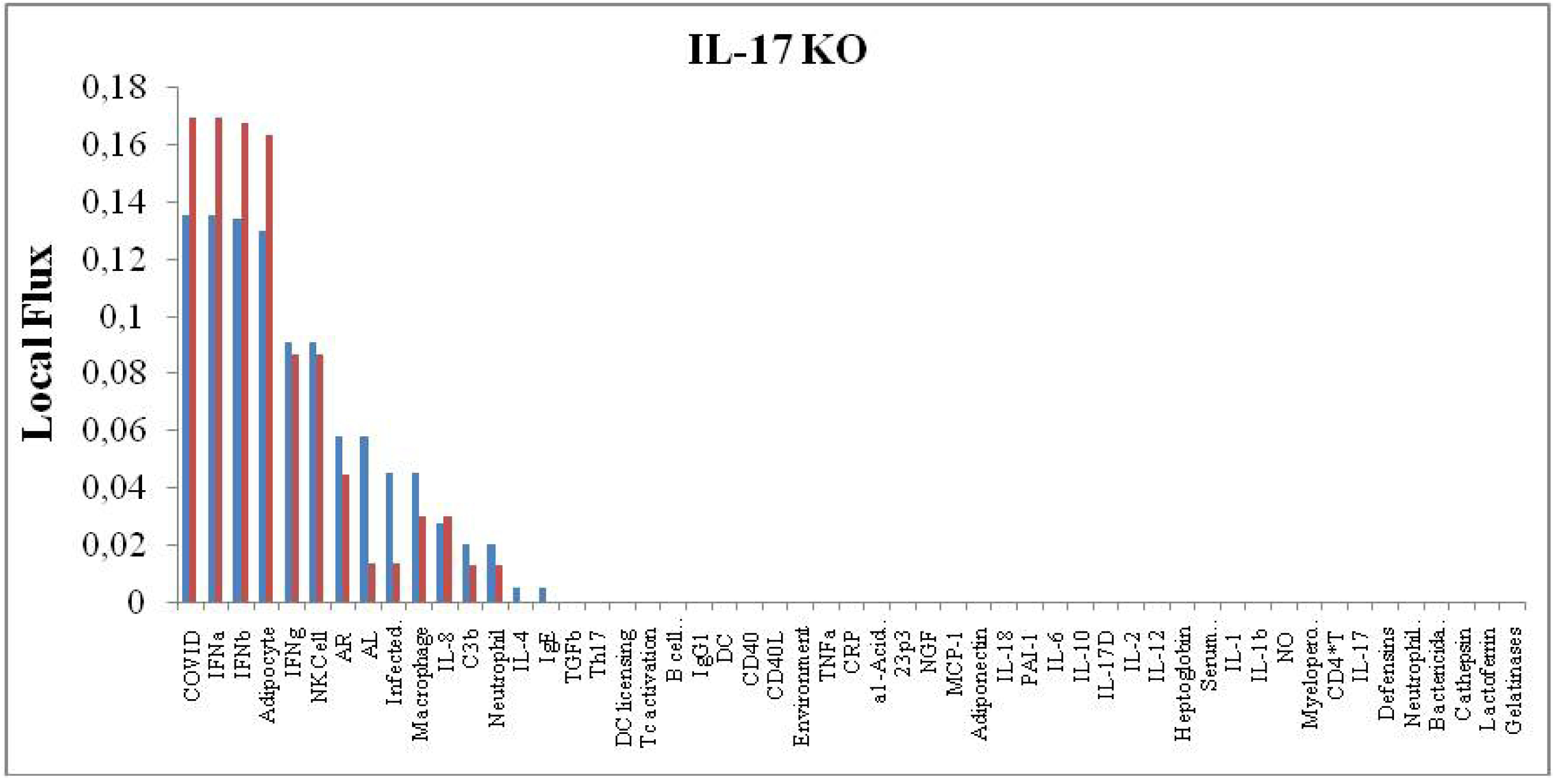
Flux Profile for IL-17 knockout and the effect on local fluxes. The blue histogram stands for the Standard Flux (without KO) and the red histogram stands for the KO flux.

**Figure 7.**
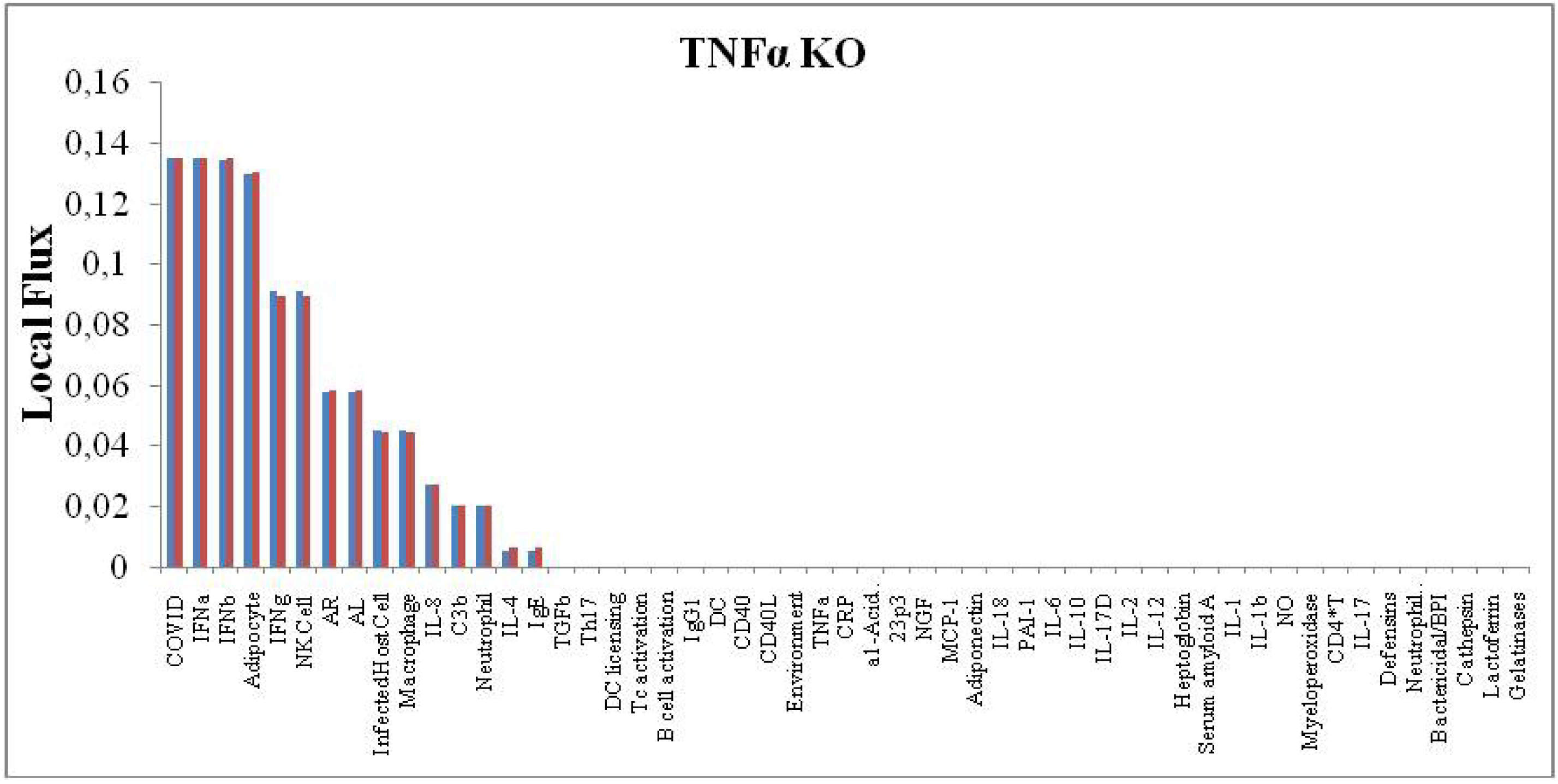
Flux Profile for TNFα knockout and the effect on local fluxes. The blue histogram stands for the Standard Flux (without KO) and the red histogram stands for the KO flux.

## 4. Discussion

The evaluation of RME in the complex network shows that the infected host cell is more important for the progression, severity and aggravation of the disease than the viral particle itself. Viruses, in general, directly or indirectly, kill or inactivate host cells, the immune response associated with this infection will cause a local inflammatory process, causing an immunopathic disease [19].

The virus-host cell relationship leads to the first sequence of immunoattractors, here, IFNs. The analysis of the KOs shows the importance of these cytokines. In all the results presented (Fig. 3-7) these molecules are of greater importance. These substances are directly related to antiviral state, decreasing cell proliferation, increasing the number of NK and CTL functions. Another important source of secretion of these cytokines is plasmocytoid DC [20].

The CD40 present on the surface of the DC cells interacts with the CD40L activating the T cells. This link avoid infection of DCs, interfere with MHC class II-mediated antigen presentation, force pMHC internalization, Tc activation and B cell activation. Tc activation feeds inflammation by stimulating IFNg, IL-10 and other cytokines. Activated B cell induces secretion of IL-10 and TGFb that induces Th17 recruitment and increased secretion of IL-17 [25,27].

Some syndrome likeX-Linked Hyper IgM andX-linked form of hyper Immunoglobulin M have a large mortality risk by respiratory infection present high alteration in CD40 mutation [28,29]. Each syndrome shows increased serum concentrationsof IgM.

The evaluation of Figure 2 shows the importance of adipose tissue as one of the main agents of interaction in this network. This tissue is a source of constant supply of TNFα, IL-1, IL-6, IL-8, IL-10, TGFb, IL-17 and IL-18. Recently, in an initial clinical study by our group, we evaluated the secretion of IL-6 and C-reactive protein, previously and after one year of Partial Duodenal Switch surgery, with removal of the greater omentum [30]. As expected, there was a decrease in these two inflammatory markers and, interestingly, patients reported decreased asthma attacks and bronchitis. Here, we observe that, based on our network results, adipose tissue is one of the main actors in the aggravation of the installed inflammatory process.

Both the infectious process by covid and the adipose tissue have secretion of large amounts of IL-8, which generate a stimulus to neutrophils. In a study by the University of Leicester, also in a network, but with a focus on proteins and receptors in this disease, they showed a robust response from neutrophils and their important participation [31].

Another interleukin, and our results, demonstrate an essential role in the severe form of a disease that manifests itself in obese people is IL-17. Our study shows this as the main one mentioned in the developed network. The role of this interleukin has been identified as associated with the risk and prognosis of Acute Respiratory Discomfort Syndrome (ARDS). Suggestion IL-17 may be a marker for risk prediction and development of ARDS [33].

Interestingly, a recent study [34] reinforced that inhibition of IL-17, which, by the way, is immunologically possible, could be a plausible strategy to prevent Acute Respiratory Distress Syndrome (ARDS) in corona virus disease 2019. Part of this statement is due to a 2017 study that evaluated the significant and important increase in IL-17 and Th17 cytokine profile in MERS-CoV [32].

## 5. Conclusions

This initial study, in a small network, pointed out the importance of chronic inflammation in the obese individual as an important factor in potentiating the disease caused by covid-19 and, in particular, the need for a clinical study focusing on IL-17. This proved to be a possible therapeutic target to minimize the potential of the disease in obese people. The expansion of the network and the association with other chronic endemic diseases have already been the subject of new studies.

## Ethics Statement

Being a mathematical experimentation work, based on data from the specific literature, the authors attest to ethics in its development.

